# *APOE ε4* gene dose effect on imaging and blood biomarkers of glial reactivity and β-amyloid pathology

**DOI:** 10.1101/2022.09.19.508484

**Authors:** Anniina Snellman, Laura L. Ekblad, Jouni Tuisku, Mikko Koivumäki, Nicholas J. Ashton, Juan Lantero-Rodriguez, Thomas K. Karikari, Semi Helin, Marco Bucci, Eliisa Löyttyniemi, Riitta Parkkola, Mira Karrasch, Michael Schöll, Henrik Zetterberg, Kaj Blennow, Juha O. Rinne

## Abstract

Increased reactivity of microglia and astrocytes is known to be present at various stages of the Alzheimer’s continuum but their relationship with core Alzheimer’s disease pathology in the preclinical stages is less clear. We investigated glial reactivity and β-amyloid pathology in cognitively unimpaired *APOE* ε4 homozygotes, heterozygotes and non-carriers using ^11^C-PK11195 PET (targeting 18-kDa translocator protein), ^11^C-PiB PET (targeting β-amyloid), brain MRI, and a preclinical cognitive composite (APCC). Plasma glial fibrillary acidic protein (GFAP) by and plasma Aβ_1-42/1-40_ were measured using single molecule array and immunoprecipitation combined with mass spectrometry, respectively. We observed that **(i)** ^11^C-PiB-binding was significantly higher in *APOE* ε4 homozygotes compared with non-carriers in all evaluated regions, **(ii)** regional ^11^C-PK11195-binding did not differ between the *APOE* ε4 gene doses or between Aβ-positive and -negative individuals, and **(iii)** higher ^11^C-PK11195-binding and plasma GFAP were associated with lower hippocampal volume, and elevated ^11^C-PiB-binding and plasma GFAP concentration with lower APCC scores. Increased glial reactivity might emerge in later stages of preclinical Alzheimer’s disease in parallel with early neurodegenerative changes.

## Introduction

The number of persons affected by Alzheimer’s disease across its pathological continuum was recently estimated to be as high as 416 million^1^. From this global estimate, 3/4 of individuals were classified as preclinical Alzheimer’s disease, characterized by the presence of beta-amyloid (Aβ) plaques but absent of clinical symptoms^1^. In addition to the hallmark pathologies, *i*.*e*., Aβ plaques and neurofibrillary tangles, inflammation in the CNS is recognized to have an important, partly independent, role in different phases of the Alzheimer’s continuum^2^. In the brain, inflammation is mainly mediated by microglia and astrocytes, which in homeostatic conditions have multiple roles in, *e*.*g*., surveillance, maintenance of the blood-brain barrier and synaptic functions^3^. In Alzheimer’s disease, compiling evidence suggests that increased microglial and astrocytic reactivity could be present during both early, possibly protective, and later, detrimental processes^4-7^.

One factor known to be closely related with both Alzheimer’s disease and CNS innate immunity responses is apolipoprotein E (apoE) that has three different isoforms, coded by the three different alleles of the *APOE* gene (*APOE* ε2, *APOE* ε3, and *APOE* ε4). The *APOE* ε4 allele is the strongest genetic risk factor of sporadic Alzheimer’s disease; it increases the risk of disease and decreases the age of onset when compared with the most common *APOE* ε3 or the protective *APOE* ε2 alleles^8^. *APOE* ε4 gene dose related increase in brain Aβ load is present already in cognitively normal individuals^9-11^, and it has been suggested to be caused by impaired degradation and clearance of Aβ, a task which is performed by glial cells and affected by apoE isoforms ^12, 13^. Most of the CNS apoE is produced by astrocytes and reactive microglia, and it has been shown to impact innate immune responses in the brain during Alzheimer’s disease pathogenesis^14-16^. In neuropathological studies, *APOE* ε4 has been seen to associate with increased microglial number in the brains of individuals with Alzheimer’s disease ^17^, and higher microglial cell reactivity around Aβ plaques in a mouse model of Aβ deposition and human *APOE* alleles^18^.

Investigation of regional glial reactivity in Alzheimer’s disease and other neurodegenerative diseases *in vivo* has been enabled by PET imaging and specific ligands such as ^11^C-PK11195 that target 18-kDa translocator protein (TSPO) as a proxy for microglial reactivity. TSPO is present in the outer mitochondrial membranes of microglia and elevated in the brain in relation to injuries or pathology^19^. In humans, increased TSPO ligand-binding has recently been suggested to represent changes in cell density rather than protein overexpression^20^, and to be mostly covered by microglia, and to a lesser extent astrocytes and endothelial cells^21, 22^. Previous studies using TSPO PET imaging have shown increased regional ligand-binding in patients with Alzheimer’s disease^23-26^, mild cognitive impairment^4, 27, 28^ and in Aβ-positive compared with -negative controls^7, 29^. However, results are partly inconclusive since also minor or no differences between diagnostic groups have been reported^30-32^.

In addition to imaging, more easily accessible biomarkers for Alzheimer’s disease pathology measured in blood have become available recently thanks to the development of more sensitive methods^33^. Soluble Aβ peptides of various lengths can be measured from plasma by combining immunoprecipitation with detection using mass spectrometry, and the plasma Aβ_1-42/1-40_ ratio has been shown to be decreased in early Alzheimer’s disease, although with lower fold changes between Aβ-positive and -negative individuals when compared to CSF Aβ_1-42/1-40_^34-36^. Unfortunately, since proteins expressed by microglia in the CNS are also present in peripheral macrophages, the development of assays targeting microgliosis is demanding, and interpretation of measurements from blood complicated^36^. However, an interesting fluid biomarker, glial fibrillary acidic protein (GFAP, a marker of reactive astrocytosis) is detectable from blood using the single molecule array (Simoa) technology, and was recently shown to be associated with Aβ deposition and increased already in early stages of Alzheimer’s disease^37-39^.

The aim of this study was to evaluate *in vivo* the differences in regional glial reactivity, Aβ deposition, and their association primarily amongst cognitively normal *APOE* ε4 homozygotes, heterozygotes and non-carriers, as well as secondary between Aβ-positive (representing Alzheimer’s pathological change or preclinical Alzheimer’s disease^40^) and Aβ-negative individuals. In addition, we aimed to investigate the association between imaging and fluid biomarkers of glial reactivity and Aβ deposition and markers of disease progression (cognitive performance and volumetric brain changes) in our cohort comprised by cognitively unimpaired participants enriched with *APOE* ε4 carrriers.

## Methods

### Study design and participants

The study design is illustrated in **Figure 1** and a detailed study protocol has previously been reported^41^. Briefly, participants in this cross-sectional, observational study, were recruited in collaboration with the local Auria biobank (Turku, Finland). Set inclusion criteria were 60-75 years of age, and CERAD total score > 62 points at screening. Main exclusion criteria were dementia or cognitive impairment; other severe neurological or psychiatric disease; diabetes; chronic inflammatory condition; and contraindication for MRI or PET imaging. The study was approved by the Ethical Committee of the Hospital District of Southwest Finland. All participants signed a written informed consent according to the Declaration of Helsinki.

**Figure 1.**
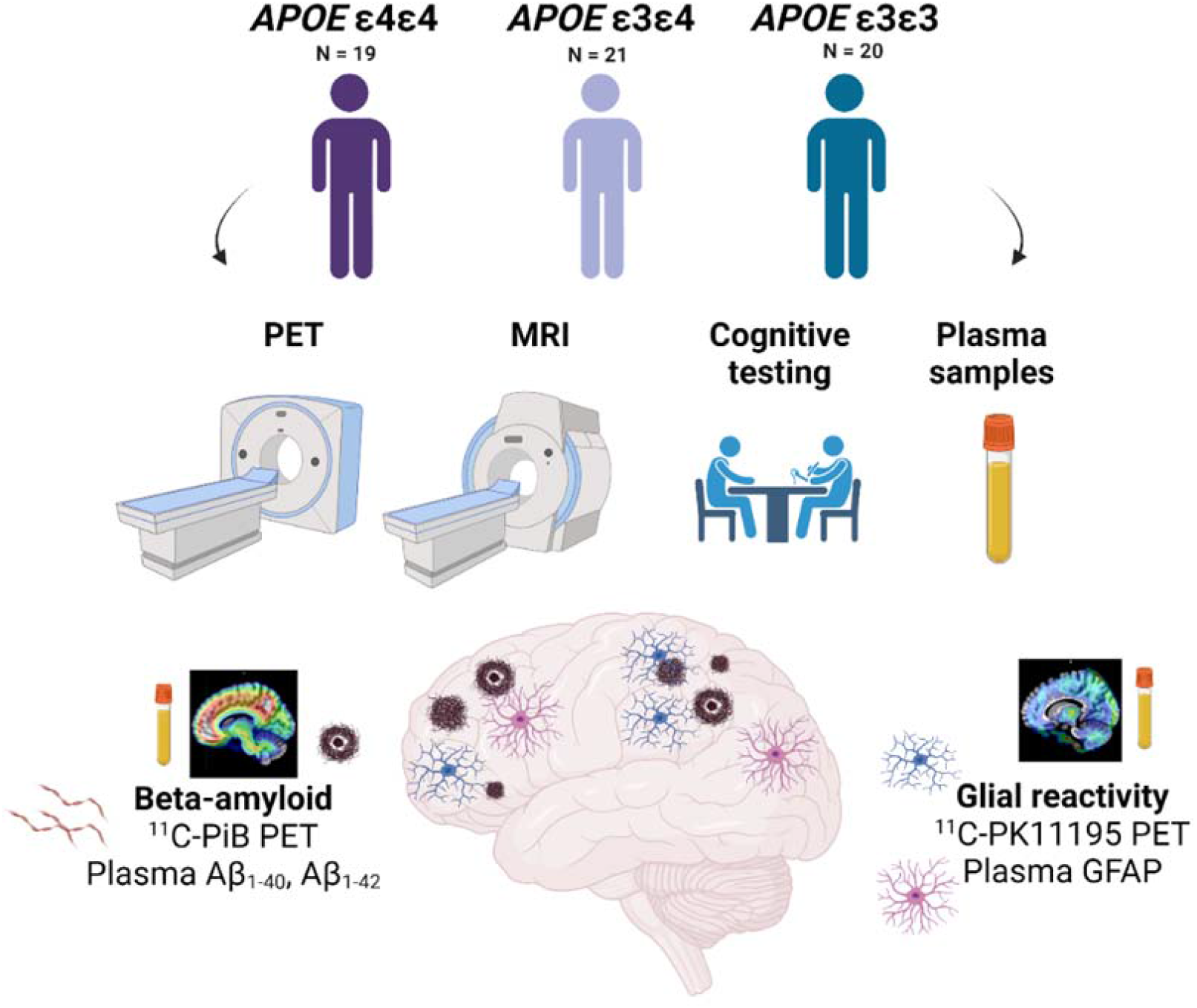
Study flowchart. Altogether 60 individuals were recruited based on their *APOE ε4* gene dose (*APOE ε4/ε4, n* = 19, *APOE ε4/ε3, n* = 21, *APOE ε3/ε3 n* = 21). All underwent positron emission tomography (PET) imaging targeting Aβ using ^11^C-PiB, 18-kDa translocator protein (TSPO) as a proxy for glial reactivity using ^11^C-PK11195, magnetic resonance imaging (MRI) and cognitive testing. A blood sample was drawn for laboratory measurements, including plasma markers of Aβ pathology (Aβ_1-40_ and Aβ_1-42_) and reactive astrocytosis (glial fibrillary acidic protein, GFAP).

### Brain imaging measurements

Structural T1-weighted brain MRI scan was performed on either a Philips Ingenuity 3.0 T TF PET/MRI (*n* = 38; Philips Healthcare, Amsterdam, the Netherlands), or a Philips Ingenia 3.0 T (*n* = 22; Philips Healthcare, Amsterdam, the Netherlands). PET scans were acquired on an ECAT high-resolution research tomograph (HRRT, Siemens Medical Solutions, Knoxville, TN). For amyloid imaging, ^11^C-PiB scans (*n* = 60) were acquired 40 to 90 minutes post injection (mean injected dose 497 (30) MBq), and for TSPO imaging, dynamic ^11^C-PK11195 scans (*n* = 57) were acquired for 60 minutes post injection (mean injected dose 494 (21) MBq). All images were reconstructed with 3D ordinary Poisson ordered subset expectation maximization algorithm (OP-OSEM3D), and list mode data was histogrammed into 8 (6 × 5 + 2 × 10min, ^11^C-PiB) and 17 (2 × 15; 3 × 30; 3 × 60; 7 × 300; 2 × 600 s, ^11^C-PK11195) time frames.

### Brain image analysis

PET and MR image preprocessing and analysis was performed using an automated pipeline at Turku PET Centre^42^ which executed the PET data frame by frame realignment, PET-MRI co-registration, FreeSurfer ROI parcellation and PET data kinetic modelling. Regional and voxel level ^11^C-PiB-binding was quantified as standardized uptake value ratios (SUVR) calculated for 60 to 90 minutes post injection, using the cerebellar cortex as reference region. Regional ^11^C-PK11195-binding was quantified as distribution volume ratios (DVR) within 20–60 min post injection using a reference tissue input Logan’s method with pseudo-reference region extracted using supervised clustering algorithm^43, 44^. Voxel-level kinetic modelling for ^11^C-PK11195 was carried out using basis function implementation of simplified reference tissue model with respect to the aforementioned clustered pseudo-reference region and with 300 basis functions calculated within the Θ_3_parameter limits 0.06 ≤ Θ_3_ ≤ 0.6.^45^ Partial volume effect (PVE)-corrected data was used for all ^11^C-PK11195 analysis in order to minimize the effect TSPO uptake in sinuses to cortical regions. PVE correction was carried out using PETPVE12 toolbox^46^ in both region-of-interest (ROI, geometric transfer matrix method) and voxel-level (Muller-Gartner method) data. ROI-level analysis for both ^11^C-PiB and ^11^C-PK11195 data was performed in *a priori* defined regions known for early Aβ deposition (prefrontal cortex, parietal cortex, anterior cingulum, posterior cingulum, precuneus, lateral temporal cortex, and a volume weighted composite containing all the regions).^41^ For ^11^C-PK11195, additional volume-weighted ROIs for transentorhinal (Braak I-II), and limbic composite (Braak III-IV) regions^47^ were analysed to investigate TSPO-binding in regions associated with early tau deposition. Details of the combined FreeSurfer regions are previously published^41^. Spatially normalized parametric SUVR and BP_ND_ images in MNI152 space were smoothed using Gaussian 8mm FWHM filter and used for all voxel-wise statistical analysis. For all figures, BP_ND_ were transformed to DVRs for clarity, using the formula: DVR = BP_ND_ + 1. Amyloid positivity was defined as cortical composite ^11^C-PiB SUVR > 1.5^48, 49^

Total hippocampal volume (left + right, ml) and total entorhinal area volume (ml) normalized for intracranial volume, age and sex were obtained from the T1-weighted MR images using an automatic cNeuro image analysis tool (Combinostics Oy, Tampere, Finland)^50, 51^. Since two different instruments were used for acquiring MRI images the used scanner was added as a covariate in all analyses including hippocampal or entorhinal volumes.

### Cognitive testing

All participants completed CERAD cognitive test battery at screening, as well as more extensive neuropsychological testing during one of the study visits as previously described.^41^ CERAD total score, mini-mental state examination (MMSE) score, and API Preclinical Cognitive Composite (APCC) score were used to investigate the association between both imaging and blood biomarkers and cognitive performance.

### Blood biomarker measurements

All plasma biomarker measurements were performed in the Clinical Neurochemistry Laboratory, Mölndal, Sweden. Plasma Aβ_1-40_ and Aβ_1-42_ concentrations were measured using an in-house immunoprecipitation mass spectrometry method (IP-MS) described in detail elsewhere^35, 52^. Briefly, Aβ peptides were immunoprecipitated from 250 μl of sample using 4G8 and 6E10 anti-Aβ antibodies (BioLegend) coupled to Dynabeads™ M-280 Sheep Anti-Mouse IgG magnetic beads and a KingFisher Flex instrument (Thermo Fisher Scientific), and further analyzed by liquid chromatography-tandem mass spectrometry (LC–MS/MS). Recombinant Aβ_1-40_ and Aβ_1-42_ peptides were used as calibrators, and heavy labelled peptides were added to both samples and calibrators for internal standards.

Plasma GFAP concentration was measured using the Single molecule array (Simoa) platform, a HD-X analyzer (Quanterix, Billerica, MA), and a commercial GFAP discovery kit (Quanterix, #102336) following the instructions provided by the manufacturer. Two internal quality control (QC) samples with mean concentrations of 100 pg/ml and 608 pg/ml were measured in the beginning and after samples in both plates. Calibrators and QC samples were measured as duplicates, and samples as singlicates. The intra-assay precision (variation within run, CV_r_ (%)) and inter-assay precision (variation between runs, CV_rw_ (%)) were < 5% and < 15%, respectively.

### Statistical analysis

All data following normal distribution are presented as mean (standard deviation, SD), otherwise as median (interquartile range, IQR). Normality of the data was established visually and from the residuals. Missing data points for each variable are presented in Supplementary methods and **Supplementary Table 1**. For continuous variables, differences in group demographics and in regional ^11^C-PiB and ^11^C-PK11195-binding between the three *APOE* ε4 gene doses were tested using one-way ANOVA with Tukey’s honest significance test, or Kruskal-Wallis test with Steel-Dwass method for multiple comparisons depending on the distribution of data. χ^2^ test was used for testing categorical variables. Associations between regional PET data and fluid biomarker concentrations were evaluated using Spearman’s rank correlation. Differences in ^11^C-PK11195-binding between amyloid positive and negative individuals were first tested with students t-test. We also wanted to see if regional ^11^C-PK11195-binding differed between amyloid positive and amyloid negative individuals accounting for *APOE* ε4 status, so we additionally tested the effect of *APOE* ε4 gene dose, amyloid positivity, and their interaction (ε4 gene dose × amyloid positivity) on ^11^C-PK11195 in *a priori* defined ROIs with linear regression models. If an interaction term with *P* < 0.1 was found, a post-hoc comparison of all groups was performed to explore the nature of the interaction.

**Table 1.** Demographics and descriptive data for cognitively unimpaired *APOE ε4* homozygotes, heterozygotes, and non-carriers included in the study.

|  | GROUP |  |  |  |
| --- | --- | --- | --- | --- |
| | *APOE* $\epsilon 4\epsilon 4$ | *APOE* $\epsilon 4\epsilon 3$ | *APOE* $\epsilon 3\epsilon 3$ | *P* |
| *n* | 19 | 21 | 20 |  |
| Age (y), mean (SD) | 67.3 (4.74) | 67.3 (4.90) | 68.3 (4.55) | 0.75 |
| Sex (M/F), *n* (%) | 7/12 (37/63) | 7/14 (33/67) | 8/12 (40/60) | 0.91 |
| Education, *n* (%) |  |  |  | 0.37 |
| **Primary school** | 7 (37) | 4 (19) | 7 (35) |  |
| **Middle or comprehensive school** | 4 (21) | 4 (19) | 3 (15) |  |
| **High school** | 7 (37) | 6 (29) | 7 (35) |  |
| **College or university** | 1 (5) | 7 (33) | 3 (15) |  |
| BMI (kg/m<sup>2</sup>), mean (SD) | 26.6 (4.48) | 26.7 (3.46) | 27.3 (4.96) | 0.86 |
| Family history of memory disorder, *n* (%) | 10 (53) | 9 (43) | 11 (55) | 0.71 |
| CERAD total score, mean (SD) | 84.4 (9.43) | 85.9 (7.98) | 86.0 (7.42) | 0.79 |
| MMSE, median (IQR) | 28 (27–29) | 29 (28–30) * | 29 (27–30) | **0.039** |
| Total leukocyte count (E9/L), mean (SD) | 5.38 (1.20) | 5.70 (1.68) | 5.22 (0.87) | 0.49 |
| <sup>11</sup>C-PIB positivity, *n* (%) | 16 (84) | 10 (48) | 8 (40) | **0.0066** |
| Computed Fazekas score, median (IQR) | 1.09 (0.98) | 0.92 (0.62) | 0.82 (0.79) | 0.8 |
| <sup>11</sup>C-PIB composite SUVR, median (IQR) | 2.13 (1.61–2.83) | 1.55 (1.43–2.02) | 1.47 (1.38–1.66) * | **0.0024** |
| <sup>11</sup>C-PK11195 composite DVR, mean (SD) | 1.34 (0.08) | 1.34 (0.05) | 1.31 (0.06) | 0.29 |

Data are presented as mean (standard deviation) or median (interquartile range) depending on the distribution.
Differences between groups were tested with one-way ANOVA with Tukey's honest significance test, or Kruskal-Wallis test with Steel-Dwass method for multiple comparisons for continuous variables. $\chi^2$ test was used for testing categorical variables. *P* value presents overall difference between groups. Significant differences in pairwise comparisons to *APOE* $\epsilon 4\epsilon 4$ homozygotes (\*) are also presented.
Abbreviations: BMI, body mass index; CERAD; Consortium to establish a Registry for Alzheimer's disease; DVR, distribution volume ratio; MMSE, mini-mental state examination; SUVR, standardized uptake value ratio.

Voxel-level differences in ^11^C-PIB and ^11^C-PK11195-binding between *APOE* ε4 gene doses were evaluated using one-way ANOVA, followed by post-hoc pairwise comparisons in Statistical Parametric Mapping (SPM12 v12; Wellcome Trust Centre for Neuroimaging, London, UK) running on MATLAB, whereas voxel-level ^11^C-PIB SUVRs and ^11^C-PK11195 BP_ND_ Spearman’s rank correlation coefficients were calculated using built-in MATLAB functions. False Discovery Rate-corrected cluster level threshold was set at *P* < 0.05. Differences in blood biomarker concentrations between *APOE* ε4 gene doses were analysed using Kruskal-Wallis test with Steel-Dwass method for multiple comparisons.

Finally, we used multivariable linear regression models adjusted for age, sex and education (and MRI scanner for models explaining hippocampal or cortical volumes) to test how well PET and fluid biomarkers of Aβ and glial reactivity could explain different cognitive and structural variables that could be interpreted as markers of disease progression. For comparison, standardized βs were calculated and presented in figures.

All statistical analyses were performed using SAS JMP Pro v.15.1.0 (SAS institute, Gary, NC) and visualizations using GraphPad Prism version 9.0.1 (GraphPad, San Diego, California, USA). A *P*-value < 0.05 (2-tailed), was considered statistically significant in all analysis, except for interaction effects, where stratified analysis was run already if *P* (interaction) < 0.1.

## Results

### Participant demographics

Demographics and descriptive data for the *APOE* ε4 gene dose groups are presented in **Table 1**. No statistically significant differences in age, sex, education, body mass index (BMI), or CERAD total score were present between the *APOE* ε4 gene dose groups (*P* > 0.37 for all). *APOE* ε4 heterozygotes had significantly higher MMSE than homozygotes (*P* = 0.036). Using a cut-off value of cortical composite ^11^C-PiB SUVR > 1.5, 84 % (*n* = 16) of the *APOE* ε4 homozygotes, 48 % (*n* = 10) of the heterozygotes, and 40.0 % (*n* = 8) of non-carriers in our cohort were classified as amyloid positive.

Age had positive correlation with ^11^C-PiB cortical composite SUVRs in *APOE* ε4 homozygotes (Rho = 0.63, *P* = 0.0039), but not in heterozygotes, non-carriers, or the whole cohort (*P* > 0.19 for all). There was no correlation between age and composite cortical ^11^C-PK11195 DVRs (*P* > 0.42 for all), plasma GFAP (*P* > 0.17 for all) or plasma Aβ_1-42/1-40_ (*P* = 0.22 for all).

For secondary analyses, we also stratified the cohort based on Aβ positivity (composite ^11^C-PiB SUVR > 1.5). Demographics are presented in **Supplementary table 2**. Significant differences between Aβ-positive and Aβ-negative individuals were found in education level (*P* = 0.046), CERAD total score (*P* = 0.0034) and MMSE score (*P* = 0.0074).

**Table 2.**
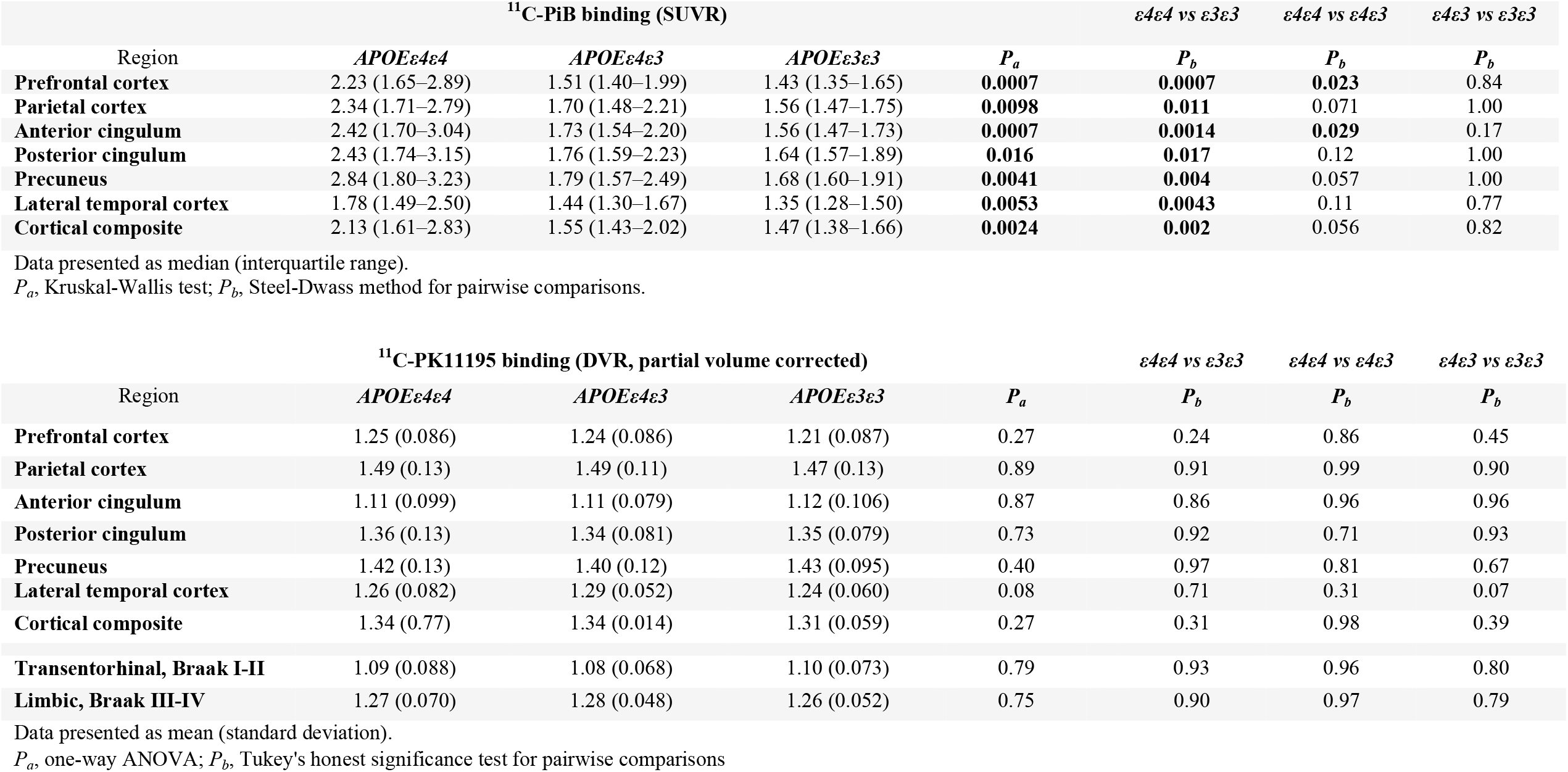
Regional ^11^C-PiB SUVR and ^11^C-PK11195 DVR values for *APOE ε4* homozygotes, heterozygotes, and non-carriers.

### Fibrillar Aβ deposition estimated by ^11^C-PiB amyloid PET

*APOE* ε4 gene dose related differences in fibrillar amyloid load were visually detectable from mean ^11^C-PiB distribution maps in regions typical for early amyloid deposition (**Figure 2A**). ROI-level analysis verified the findings, revealing significant differences in ^11^C-PiB-binding between gene doses in all evaluated regions (*P* < 0.016 for all regions, Kruskal-Wallis test). After post-hoc comparison of all groups, ^11^C-PiB-binding was significantly higher in *APOE* ε4 homozygotes compared with heterozygotes in the anterior cingulum (*P* = 0.029) and prefrontal cortex (*P* = 0.023), and in all evaluated regions when compared with non-carriers (*P* < 0.017 for all regions) (**Figure 2B, Table 2**).

**Figure 2.**
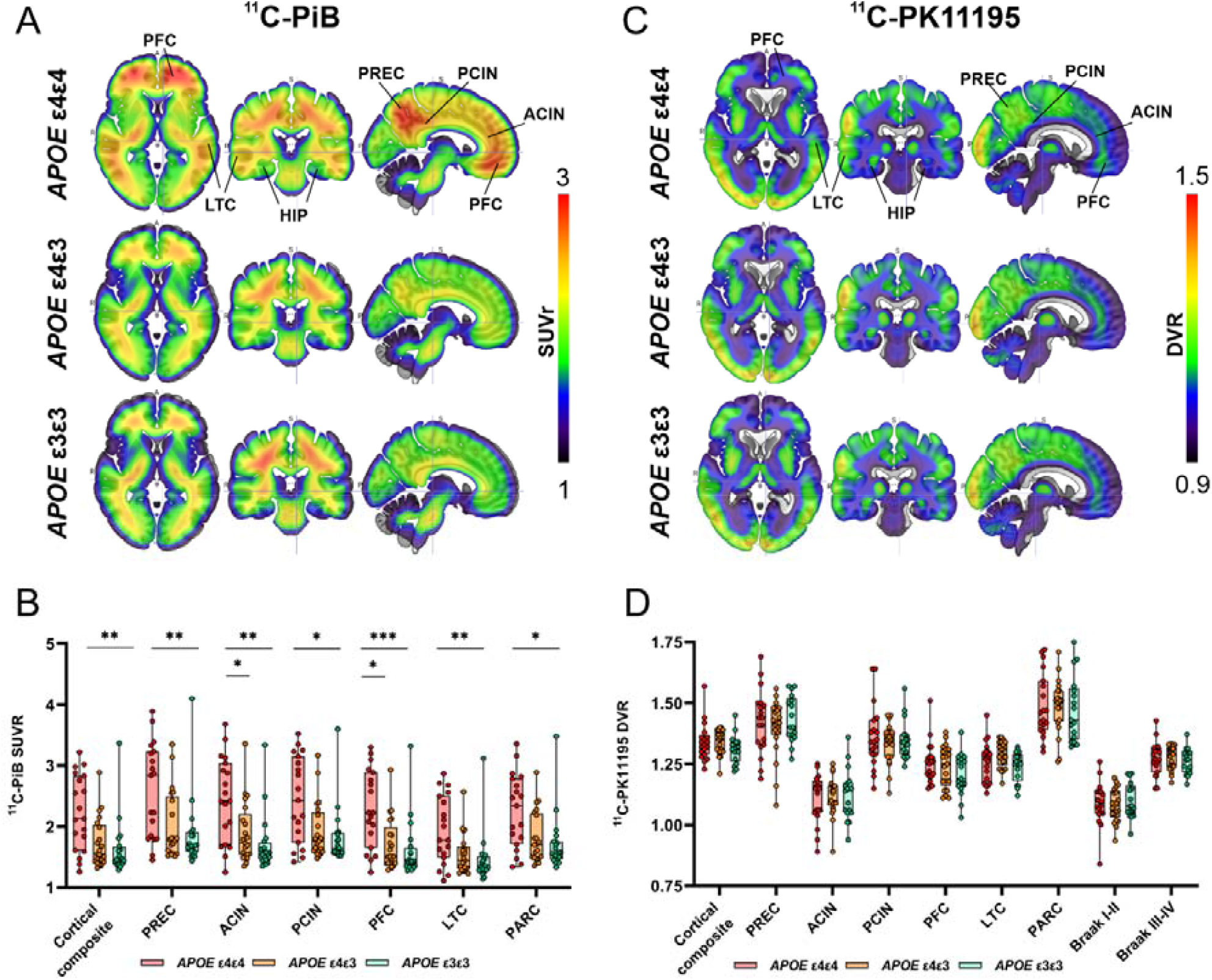
Mean ^11^C-PiB and ^11^C-PK11195 distribution maps and regional ligand-binding in cognitively unimpaired volunteers stratified by *APOE* ε4 gene dose. **(A)** Mean ^11^C-PiB standardized uptake value ratio (SUVR) distribution maps and **(B)** region-of-interest analysis showed significantly higher uptake in *APOE ε4* homozygotes compared with non-carriers in all evaluated regions and compared with heterozygotes in anterior cingulate (ACIN) and prefrontal cortex (PFC) (Kruskal-Wallis test with Steel-Dwass method for multiple comparisons). **(C)** Mean ^11^C-PK11195 standardized distribution volume ratio (DVR) maps showed regional differences in tracer-binding **(D)** but no significant differences between the *APOE ε4* gene dose groups (One-way ANOVA with Tukey’s honest significance test for multiple comparisons). HIP, hippocampus; PARC, parietal cortex; PCIN, posterior cingulate cortex; PREC, precuneus.). * *P* < 0.05; ** *P* < 0.01; *P* < 0.001

Voxel-level comparisons verified the findings showing significantly higher ^11^C-PiB-binding in the prefrontal cortex, precuneus and lateral temporal cortex of the *APOE* ε4 homozygotes compared with non-carriers (**Supplementary Figure 1A**). Weaker effects with similar spatial distribution were seen in *APOE* ε4 homozygotes compared with heterozygotes (**Supplementary Figure 1B**). No significant clusters were found when comparing heterozygotes and non-carriers.

### Regional TSPO-binding estimated by ^11^C-PK11195 PET

Mean ^11^C-PK11195 DVR distribution maps for each *APOE* ε*4* gene dose are shown in **Figure 2C**. In contrast to the significant differences in fibrillar amyloid load measured by amyloid PET, we did not observe any differences in TSPO-binding between *APOE* ε*4* gene doses (*P* > 0.08 for all, one-way ANOVA, **Figure 2D, Table 2)** measured by ^11^C-PK11195 PET. In agreement with the ROI-level analyses, no significant clusters were detected in voxel-level comparisons between the *APOE* ε4 gene dose groups.

For secondary analysis, we stratified the cohort based on Aβ-positivity (^11^C-PiB SUVR > 1.5). Similar to the analyses stratified by *APOE* ε*4* gene dose, we found no regional differences in TSPO-binding between Aβ-positive and -negative individuals (*P* > 0.21 for all regions, Student’s t test, **Supplementary Table 3**). To further evaluate the possible effects of amyloid status on TSPO-binding in different *APOE* ε*4* gene doses, we analyzed also the interaction of Aβ-positivity × *APOE* ε*4* gene dose for predicting regional TSPO-binding. Whereas amyloid status (accounted for *APOE* ε*4* gene dose) did not have a significant effect on TSPO-binding (*P* > 0.28 for all regions), the interaction term approached statistical significance in the cortical composite (*P* = 0.090), lateral temporal cortex (*P* = 0.063), transentorhinal (Braak I-II, *P* = 0.052), and limbic (Braak III-IV, *P* = 0.019) ROIs (**Table 3**). In those regions, median TSPO-binding was higher in amyloid positive *APOE* ε*4* carriers than in non-carriers, and, interestingly, also in Aβ-negative non-carriers compared with Aβ-positive non-carriers (**Supplementary Figure 2**). However, these differences did not reach statistical significance after post-hoc comparison between all six groups.

**Table 3.** Test effects from multivariate linear regression models explaining regional ^11^C-PK11195 binding.

| | *APOE* $\epsilon 4$ gene dose | | Aβ status | | *APOE* $\epsilon 4$ gene dose × Aβ status | |
| --- | --- | --- | --- | --- | --- | --- |
|  | F Statistic | *P* | F Statistic | *P* | F Statistic | *P* |
| Anterior cingulum | 0.034 | 0.97 | 1.21 | 0.28 | 0.0011 | 1.00 |
| Posterior cingulum | 0.10 | 0.90 | 0.001 | 0.97 | 2.11 | 0.13 |
| Lateral temporal cortex | **3.74** | **0.031** | 0.062 | 0.80 | 2.93 | 0.062 |
| Parietal cortex | 0.14 | 0.87 | 0.0053 | 0.94 | 1.75 | 0.18 |
| Prefrontal cortex | 1.14 | 0.33 | 0.0046 | 0.95 | 0.91 | 0.41 |
| Precuneus | 0.71 | 0.71 | 0.22 | 0.64 | 0.68 | 0.51 |
| Cortical composite | 1.05 | 0.36 | 0.021 | 0.91 | 2.53 | 0.09 |
| Transentorhinal, Braak I-II | 0.67 | 0.52 | 0.23 | 0.64 | **3.14** | **0.052** |
| Limbic, Braak III-IV | 1.05 | 0.36 | 0.035 | 0.85 | **4.28** | **0.019** |

### Correlation between ^11^C-PiB and ^11^C-PK11195-binding

No significant correlation between ^11^C-PiB and PVE-corrected ^11^C-PK11195-binding was present in any of the *a priori* chosen ROIs in the total study population (Rho = −0.11-0.12, *P >* 0.35 for all, **Supplementary Table 4**). However, when stratified by *APOE* ε4 gene dose, higher composite ^11^C-PiB-binding associated with higher TSPO-binding in the cortical (Rho = 0.46, *P* = 0.043), and limbic (Rho = 0.49, *P* = 0.032) composite ROIs in *APOE* ε4 homozygotes (**Figure 3A**), but not in *APOE* ε4 heterozygotes. In contrast, a negative correlation was observed for non-carriers in the transentorhinal (Rho = −0.63, P = 0.0065) and limbic ROIs (Rho = −0.68, *P* = 0.0025).

**Figure 3.**
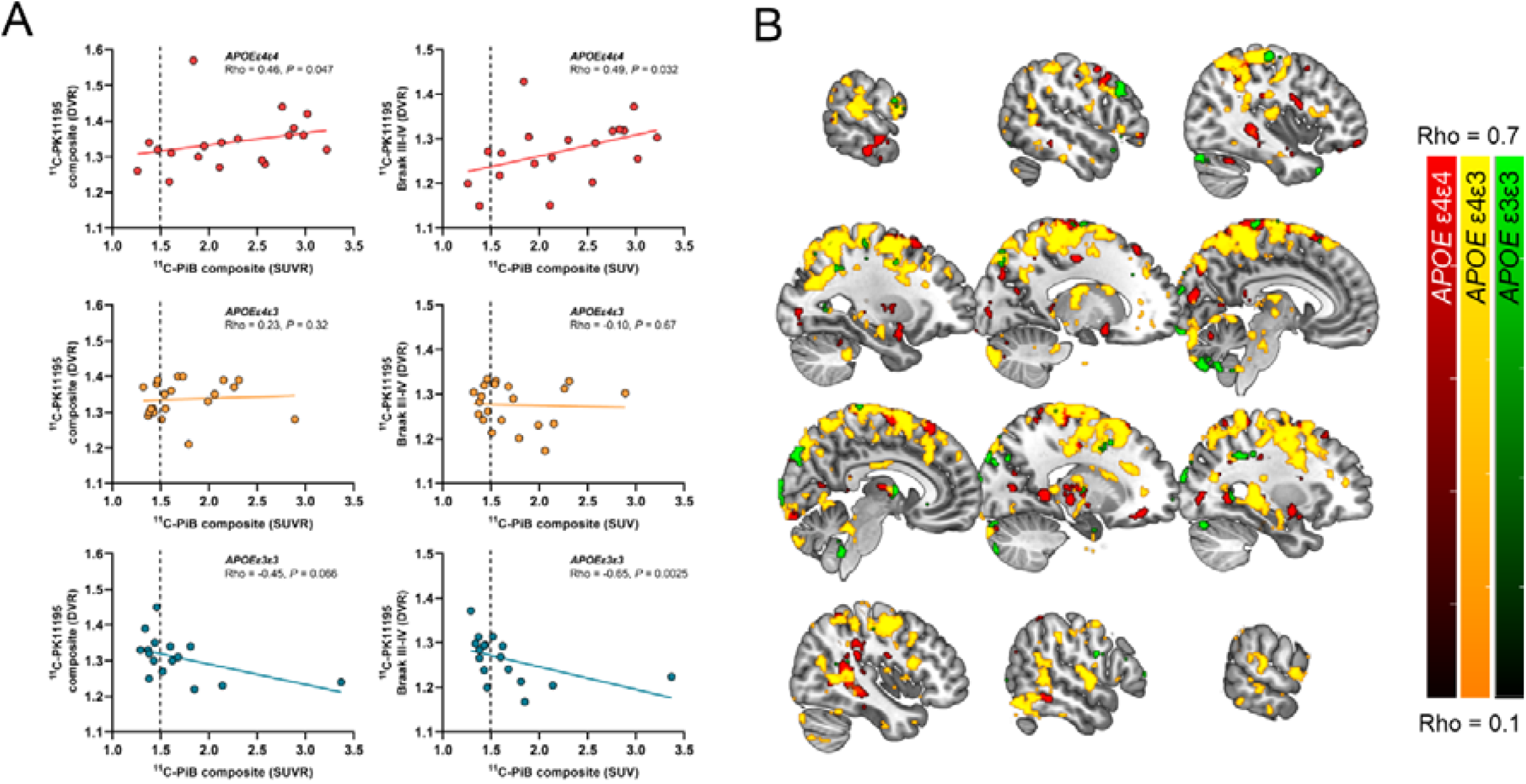
Regional association between amyloid PET and TSPO PET in cognitively unimpaired volunteers stratified by *APOE* ε4 gene dose. **(A)** Scatterplots from ROI level data showed positive correlation (Spearman’s rank correlation) for *APOE* ε4 carriers in cortical and Braak III-IV composite regions, whereas negative associations were present for non-carriers. **(B)** Most significant voxel-wise positive correlations between ^11^C-PiB and ^11^C-PK11195-binding were present in the *APOE* ε4/ε3 heterozygotes (yellow scale) and in *APOE* ε4 homozygotes (red scale), whereas only sparse significant voxels were seen in non-carriers (green scale). Partial volume corrected ^11^C-PiB SUVR and ^11^C-PK11195 BP_ND_ images smoothed using Gaussian 8mm FWHM filter were used for all voxel-wise analysis. False Discovery Rate corrected cluster level threshold was set at *P* < 0.05.

Voxel-wise analysis (not limited to specific predefined regions) did reveal clusters with significant correlation between ^11^C-PiB- and ^11^C-PK11195-binding in both *APOE* ε4 homozygotes (**Figure 3B**, red scale) and heterozygotes (**Figure 3B**, yellow scale), whereas only small spare clusters were found in non-carriers (**Figure 3B**, green scale). However, many of the clusters were located outside our primary regions of interest (chosen based on presence of early amyloid or tau pathology), such as in the white matter and the paracentral lobule.

### Astroglial reactivity estimated by plasma GFAP

Absolute plasma GFAP concentrations were higher in *APOE* ε4 homozygotes (186 pg/ml, 124-269) compared with *APOE* ε4 heterozygotes (150 pg/ml, 104-170) and non-carriers (128 pg/ml, 105-147), (*P* = 0.077, Kruskal-Wallis test, **Figure 4A**). A trend towards positive association between plasma GFAP and cortical ^11^C-PiB-binding was present in the whole cohort (Rho = 0.23, *P* = 0.085), and a significant positive correlation was observed in Aβ-positive individuals (Rho = 0.34, *P* = 0.048). No association between plasma GFAP and cortical TSPO-binding was present in the whole cohort (Rho = 0.064, *P* = 0.64) or in Aβ-positive individuals (Rho = 0.13, *P* = 0.47, **Figure 4A)**.

**Figure 4.**
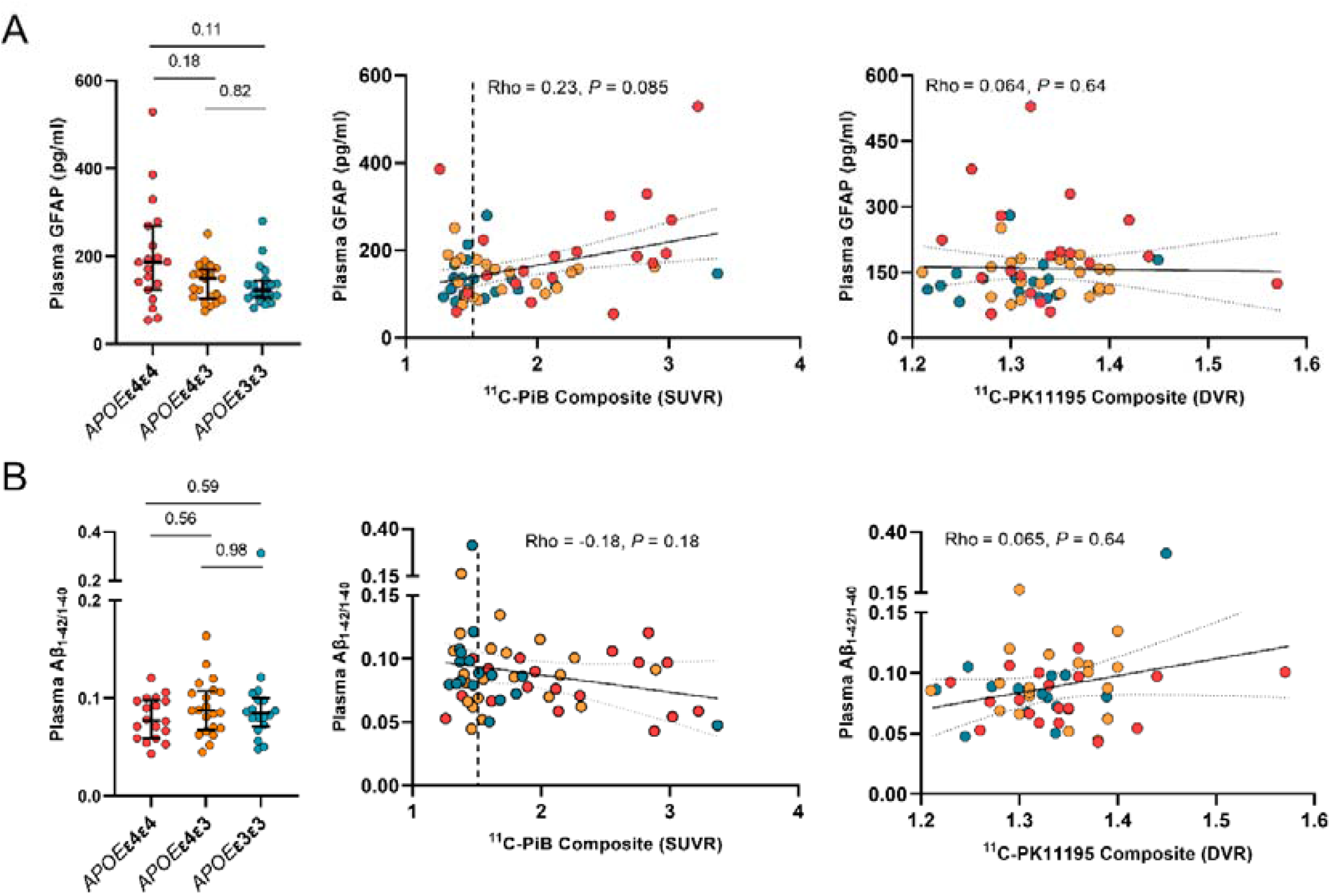
Plasma GFAP and plasma Aβ_1-42/1-40_ concentrations in cognitively unimpaired volunteers stratified by *APOE* ε4 gene dose. Differences in biomarker concentrations between *APOE* ε4 gene doses, correlations with cortical composite amyloid PET standardized uptake value ratios (SUVRs), and TSPO PET distribution volume ratios (DVRs) for (**A)** plasma glial fibrillary acidic protein (GFAP) and (**B)** plasma Aβ_1-42/1-40_. Differences between groups were tested with Kruskal-Wallis with Steel-Dwass method for multiple comparisons, and correlations with Spearman’s rank correlation.

### Soluble Aβ concentrations estimated by plasma Aβ_1-42/1-40_

Despite the clear differences in regional Aβ PET, plasma Aβ_1-42/1-40_ was not significantly different between *APOE* ε4 homozygotes (0.077, 0.059-0.098), *APOE* ε4 heterozygotes (0.087, 0.068-0.11)), and non-carriers (0.086, 0.076-0.10) (*P* = 0.50, Kruskal-Wallis test, **Figure 4B**). In our cohort, plasma Aβ_1-42/1-40_ did not correlate with either cortical composite amyloid load measured by ^11^C-PiB PET (Rho = −0.18, *P* = 0.18), or with cortical composite TSPO-binding measured by ^11^C-PK11195-binding (Rho = 0.065, *P* = 0.64; **Figure 4B**).

### Biomarker associations with cognitive performance, hippocampal and cortical volume: markers for disease progression

Finally, we wanted to compare how the different biomarkers associate with cognitive (MMSE, CERAD total score, APCC score) and structural variables (total hippocampal and entorhinal volume) that could be seen as proxies for future disease progression (**Figure 5**). In the whole cognitively unimpaired cohort, higher cortical composite ^11^C-PiB-binding (β_std_ = −0.29 (95% CI −0.52 to −0.067), *P* = 0.012), but not higher ^11^C-PK11195-binding (β_std_ = −0.045 (−0.26 to 0.17), *P* = 0.68), was associated with lower APCC scores. However, higher cortical ^11^C-PK11195-binding was associated both with lower hippocampal volume (β_std_ = −0.36 (−0.61 to −0.12), *P* = 0.0047) and entorhinal volume (β_std_ = −0.47 (−0.72 to −0.22), *P* = 0.0004). Higher plasma GFAP concentration was associated with both lower hippocampal volume (β_std_ = −0.35 (−0.61 to −0.086), *P* = 0.010), MMSE (β_std_ = −0.35 (−0.59 to −1.10), *P* = 0.0060) and APCC scores (β_std_ = −0.29 (−0.51 to −0.070), *P* = 0.011). Plasma Aβ_1-42/1-40_ was not associated with any of the cognitive or volumetric variables (P > 0.18 for all analysis). All models were adjusted for age, sex and education.

**Figure 5.**
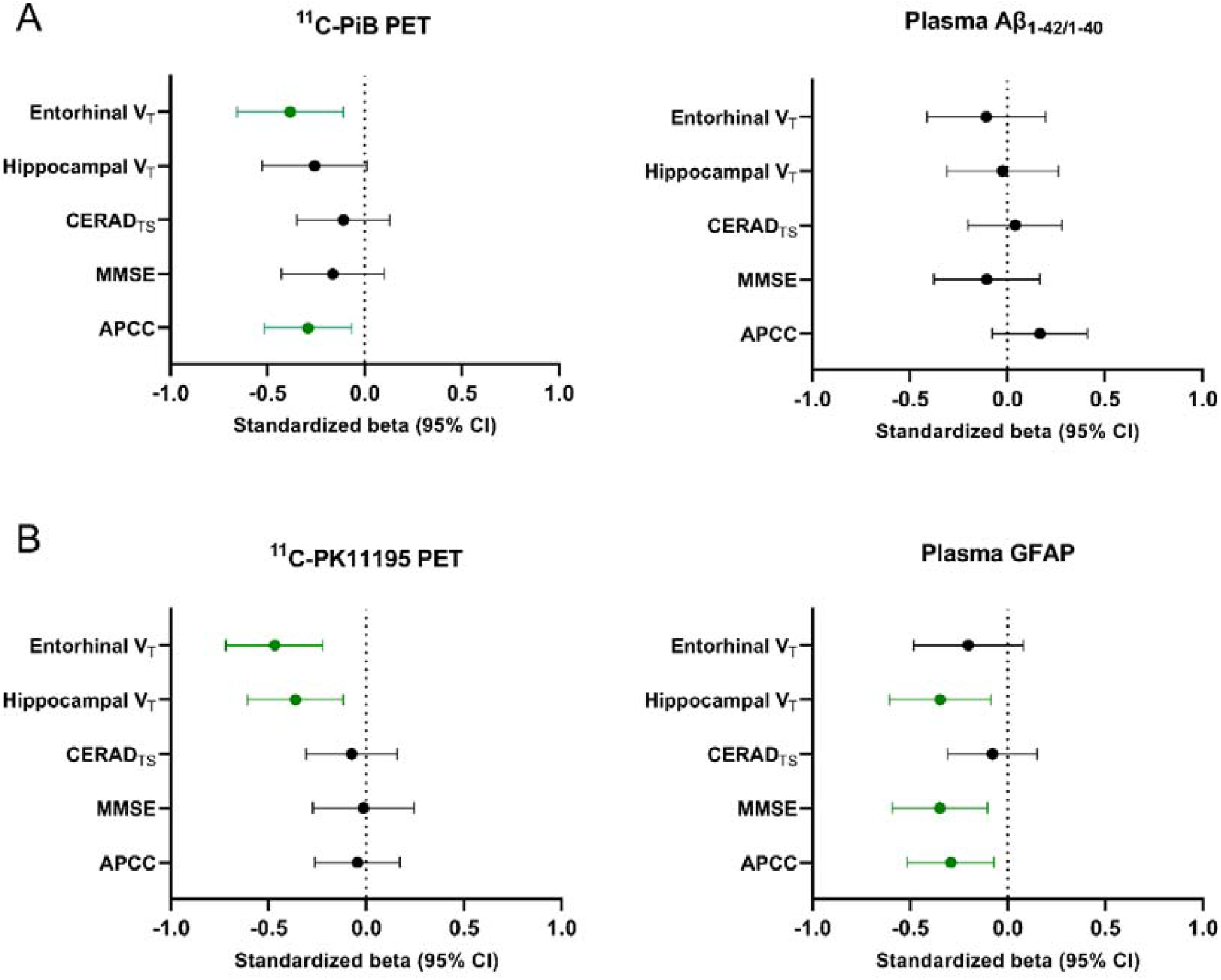
Comparison of PET and blood biomarkers of Aβ deposition and glial reactivity and their association with cognitive performance and brain structure. **(A)** Higher cortical composite ^11^C-PiB-binding, but not plasma Aβ_1-42/1-40_, was associated with lower entorhinal volumes and lower scores in the Alzheimer’s Prevention Initiatives preclinical cognitive composite (APCC) battery. (**B)** Cortical composite ^11^C-PK11195 PET was associated only with lower hippocampal and entorhinal volume, whereas elevated plasma GFAP levels were associated with lower APCC and Mini Mental State Examination (MMSE) scores. The results are shown as standardized estimates (betas) derived from liner models adjusted for age, sex, and education (and used MRI scanner for structural variables). CERAD_TS_, Consortium to Establish a Registry for Alzheimer’s Disease total score; V_T_, total volume

## Discussion

Microglial, and recently also astrocytic reactivity, have been suggested to be early events possibly present in a bi-phasic fashion during the long Alzheimer’s disease continuum.^3, 4, 6^ In previous human *in vivo* studies, increased TSPO ligand-binding has been reported to be present already in amyloid-positive mild cognitive impairment and amyloid positive controls.^7, 24, 27, 29^ Thus, we hypothesized that if such early changes are present, they should be detected in either cognitively normal *APOE* ε4 homozygotes or *APOE* ε4 heterozygotes, both representing a genetically increased risk for Aβ accumulation and sporadic Alzheimer’s disease.

First, we demonstrated that Aβ deposition in the brain increased in an *APOE* ε4 gene dose dependent fashion; significantly elevated cortical ^11^C-PiB retention was present in *APOE* ε4 homozygotes compared with both heterozygotes and non-carriers in all evaluated regions. These findings are in line with previous PET studies,^9-11, 53, 54^ as well as with a recent study by the Amyloid Biomarker Study Group summarizing *APOE* ε4 gene dose related effects on temporal course of Aβ accumulation.^55^ Similar to a previous *APOE* ε4 gene dose study,^9 11^C-PiB-binding correlated with age only in *APOE* ε4 homozygotes and in our cohort, all *APOE* ε4/ε4 participants over the age of 63 were already Aβ-positive. Contrary to our expectations, significant differences in ^11^C-PiB-binding between *APOE* ε4 heterozygotes and non-carriers were not detected. Approximately 50% of the heterozygotes included in our study were still Aβ-negative, whereas 40% of non-carriers were classified as Aβ-positive. We also had two highly ^11^C-PiB-positive non-carriers (with cortical composite SUVRs of 3.4 and 2.2) without known risk factors included in our cohort. Interestingly, both of these individuals had lower cortical TSPO-binding and plasma GFAP levels than *APOE* ε4 homozygotes with similar Aβ load quantified by PET (Figure 3A and 4A).

Despite the clear differences in fibrillar Aβ load, we did not find significant regional differences in TSPO-binding among cognitively normal individuals with different *APOE* ε4 gene dose, or between cognitively normal Aβ-positive (presenting Alzheimer’s pathological change or preclinical Alzheimer’s disease) and Aβ-negative individuals. Previously, most robust increases in TSPO-binding have been found in Alzheimer’s dementia in comparison to controls,^23-25, 56^ but also in Aβ-positive MCI.^4, 27, 28^ In addition, using second generation TSPO ligands ^18^F-DPA-714 and ^11^C-PBR28, increased TSPO-binding has been reported between Aβ-positive and -negative controls,^7, 29^ whereas another study using another second generation TSPO ligand, ^18^F-FEPPA, reported no differences in regional TSPO-binding between amnestic MCI patients and healthy volunteers.^32^ Our findings are in line with Knezevic and colleagues, since despite clearly increased fibrillar Aβ load, we were not able to replicate the reported increased TSPO-binding in Aβ-positive “at-risk” individuals using ^11^C-PK11195 PET, even with a larger sample size compared with previous reports. Our study included approximately 20 participants in each *APOE* ε4 gene dose group, and 34 cognitively unimpaired Aβ-positive individuals, whereas the previous studies included only six^7^ or seven^29^ Aβ-positive controls. These differences between studies are likely explained by the highly dynamic nature of inflammatory processes in health and disease; the rather low number of subjects (especially of individuals with prodromal or preclinical Alzheimer’s disease) included in neuroimaging studies; and the known limitations of the TSPO method regarding its specificity. It should also be noted that our cohort, and especially its Aβ-positive participants, are highly enriched with *APOE* ε4 carriers. Since *APOE* is suggested to be also directly linked to immune responses and activation state of microglia in Alzheimer’s disease,^14, 57, 58^ we cannot exclude a direct negative effect of *APOE* ε4 to microglial response-related Aβ pathology that could explain the lack of increased TSPO-binding in Aβ-positive participants in our cohort.

Activated microglia is known to be located in the proximity of Aβ plaques in Alzheimer’s disease, and using PET imaging *in vivo*, Aβ pathology has been shown to correlate with TSPO-binding in some, although not all studies.^7, 28, 30, 59, 60^ In our partial volume corrected ROI level analysis, cortical composite TSPO-binding was moderately correlated with Aβ PET signal only in *APOE* ε4 homozygotes. In a previous study using a second generation TSPO ligand (^11^C-PBR28) and including cognitively normal elderly individuals and participants with mild cognitive impairment, TSPO-binding was associated with increased Aβ PET signal only in Aβ-negative individuals.^61^ In another study, correlations were stronger in MCI compared with Alzheimer’s disease.^59^ In agreement, our voxel level analysis showed significant correlations both in cognitively normal *APOE* ε4 homozygotes and heterozygotes. However, significant clusters were found also in regions outside our *a priori* chosen regions of interest, such as the white matter, suggesting that these effects might not all be related to Alzheimer’s pathological change.

Third, Aβ positivity modulated the effect of *APOE* ε4 gene dose on ^11^C-PK11195-binding in regions known for early tau deposition, and a trend towards elevated TSPO-binding was present in Aβ-positive *APOE* ε4 carriers compared with Aβ-positive non-carriers. In addition to Aβ, *APOE* ε4 is known to accelerate tau pathology, that again has been suggested to be closely associated with microglial reactivity,^62^ and increased tau PET signal in the entorhinal cortex has been reported for cognitively unimpaired Aβ-positive *APOE* ε4 homozygotes and heterozygotes compared with Aβ-positive non-carriers.^53^ Since Aβ build up starts earlier in *APOE* ε4 carriers, we could hypothesize that increased tau deposition in *APOE* ε4 carriers would be driving this interaction. Unfortunately, lack of tau PET or CSF tau measurements in our cohort prevented us from investigating the interaction with TSPO-binding and tau further in our cohort.

During recent years, significant efforts have been made to measure various biomarkers of Alzheimer’s disease pathology in plasma that would provide a less invasive and more easily accessible alternative to brain imaging and lumbar punction.^33^ Here, despite clear differences in fibrillar Aβ levels measured by PET, we did not see significant differences between the *APOE* ε4 gene doses in plasma Aβ_1-42/1-40_ measured by previously described IP-MS method.^35^ Plasma Aβ_1-42/1-40_ was previously reported to correlate with global cortical Aβ PET signal in another study including cognitively normal individuals using the same IP-MS method.^52^ We could not replicate this finding in our cohort, comprised of slightly older, and highly *APOE* ε4-enriched cognitively normal participants; although a trend towards negative association could be seen in the whole cohort. Plasma GFAP has been recently reported to be an early marker of Alzheimer’s disease pathology, that strongly correlates with Aβ pathology,^39, 63^ but not with tau when accounting for Aβ.^37^ In our cohort, plasma GFAP levels showed elevated concentrations in the most Aβ positive individuals and correlated with composite amyloid PET SUVRs. Interestingly, plasma GFAP was the only biomarker showing significant associations with both cognitive performance and entorhinal and hippocampal volumes, that could be considered as markers for progression in the Alzheimer’s continuum. Plasma GFAP concentration did not correlate with composite TSPO-binding (Figure 4). This is not surprising, considering that plasma GFAP is expected to reflect more astrocytic reactivity associated with Aβ pathology,^39^ whereas TSPO PET is thought to reflect microglial density.^20^ Our results with GFAP support the previous findings suggesting that reactive astrocytosis is present already in cognitively normal individuals and related to Aβ pathology.^37, 39, 63^

Last, we also wanted to compare all the biomarkers and their associations with cognitive and structural variables that could serve as proxies for disease progression in our “at-risk” cohort. We found a negative association between composite cortical TSPO-binding and hippocampal and entorhinal volumes, suggesting that more global elevation in TSPO-binding, and thus microglial density, could be present in individuals with subtle neurodegeneration. Interestingly, higher plasma GFAP associated with both lower cognitive performance and lower hippocampal volume in our cognitively normal cohort. Previously, Hamelin and colleagues reported a positive correlation with both hippocampal volume and MMSE score, suggesting that higher glial reactivity associated with higher hippocampal volume would likely be protective.^7^ However, our study population is composed of only cognitively unimpaired individuals highly enriched for *APOE* ε4 carriers, and all having MMSE scores > 25, thus likely presenting more subtle structural brain changes compared with the population of the previous study. In addition, we did not find any association with TSPO-binding and MMSE, CERAD total score, or the preclinical cognitive composite, in line with other studies performed with ^11^C-PK11195.^28^ Based on our results, increased TSPO-binding in the preclinical phase, at least in *APOE* ε4 carriers, could be more related to a later preclinical phase when subtle neurodegeneration already starts to be present.

The strength of this study is our well characterized and balanced cohort of cognitively unimpaired participants stratified by their *APOE* ε4 gene dose, and a relatively large number of rare homozygotic carriers of the *APOE* ε4 allele. However, this study does not go without limitations. First, we were not able to include tau PET or CSF tau measurement. Second, even though ^11^C-PK11195 has shown robust changes in primary inflammatory conditions such as multiple sclerosis, it has been suggested that its sensitivity is limited and outperformed by the second generation TSPO ligands, such as ^11^C-PBR28. However, affinity of the second generation TSPO ligands is affected by a single nucleotide polymorphism rs6971 in the TSPO gene, leading to division of people into high, mixed, and low affinity binders. Due to the difficulty of recruiting rare homozygotic *APOE* ε4 carriers, we wanted to avoid the unfortunate scenario of having multiple homozygotic participants excluded due to low-binding TSPO genotype.

In conclusion, our study on cognitively unimpaired “at-risk” individuals carrying either one or two copies of the *APOE* ε4 gene showed clear differences in fibrillar Aβ load in the brain, but the changes were not accompanied by increased glial reactivity as measured with TSPO PET either in *APOE* ε4 carriers, or in Aβ-positive individuals, presenting preclinical Alzheimer’s disease. Plasma GFAP concentration associated with Aβ deposition in Aβ-positive individuals only. These findings suggest that in cognitively unimpaired *APOE* ε4 carriers, neuroinflammatory processes measured by TSPO PET are not closely related to Aβ accumulation, but rather to a more advanced preclinical phase of Alzheimer’s disease where Aβ accumulation is accompanied by subtle structural changes.

## Acknowledgements

The participants of ASIC-E4 study are warmly acknowledged for their commitment to the study during these challenging times. The authors would also like to acknowledge the staff of Turku PET Centre, and Auria biobank for their assistance during the recruitment and data collection for this study. We also want to thank Dr Tomi Karjalainen for all help with the automated analysis pipeline.

## Abbreviations

Aβ: beta-amyloid peptide
*APOE*: Apolipoprotein E gene
ApoE: Apolipoprotein E
TSPO: 18-kDa translocator protein
GFAP: Glial fibrillary acidic protein
GFAP: glial fibrillary acidic protein
TSPO: 18-kDa translocator protein

